# Serum-dependent recruitment of the chromatin remodeler CHD8 to promoters is mediated by the ERK-ELK pathway

**DOI:** 10.1101/2022.09.09.507301

**Authors:** Alicia Subtil-Rodríguez, Elena Vázquez-Chávez, Jose A. Guerrero-Martínez, María Ceballos-Chávez, José C. Reyes

## Abstract

Chromodomain helicase DNA binding protein 8 (CHD8) is a chromatin remodeler of the SNF2 family involved in gene transcription regulation. It has been shown that CHD8 is required for cell proliferation, cell differentiation and central nervous system development. In fact, CHD8 haploinsufficiency causes a human syndrome characterized by autism, macrocephaly, gastrointestinal complaints and some other clinical characteristics. However, the mechanism by which CHD8 controls transcription and how it is recruited to its targets in the chromatin is still unclear. We have previously shown that serum depletion causes that CHD8 detaches from chromatin. Here we demonstrate that serum-dependent recruitment of CHD8 to promoters requires the extracellular signal-regulated kinase (ERK)/ ETS-like (ELK) branch of the mitogen-activated protein kinase (MAPK) pathway. Our analysis of genomic occupancy data shows that CHD8 binding sites were strongly enriched in ELK1 and ELK4 DNA binding motifs and that CHD8 and ELK1 co-occupy multiple transcription start sites. We show that ELK1 and ELK4 are required for normal recruitment of CHD8 to the promoters of *CCNA2, CDC6, CCNE2, BRCA2* and *MYC* genes. However, CHD8 is dispensable for ELK1 and ELK4 binding. Genome wide transcriptomic analysis evidenced that serum-dependent activation of a subset of immediate early genes, including the well-known ELK1 target gene *FOS*, was impaired upon depletion of CHD8. In summary, our results uncover the role of the ERK/ELK pathway in CHD8 recruitment to chromatin and provide evidences indicating a role of CHD8 in regulating serum-dependent transcription.

## Introduction

Chromodomain helicase DNA binding protein 8 (CHD8) is a chromatin remodeler of the SNF2 family able of reshaping the structure of a nucleosome in vitro, through the hydrolysis of ATP (Thompson et al., 2008). The inactivation of the *Chd8* gene by homologous recombination in mice is lethal in homozygosis, causing the embryo to stop its development in stages prior to implantation (Nishiyama et al., 2004). Genomic studies showed that CHD8 binds to transcriptionally active promoters enriched in histone H3 dimethylated or trimethylated in lysine 4 (H3K4me2, H3K4me3) (Ceballos-Chavez et al., 2015; Cotney et al., 2015; Subtil-Rodriguez et al., 2014; Sugathan et al., 2014) (reviewed in (Wade et al., 2018)). Gene targets of CHD8 often encode transcriptional or chromatin regulators and proteins involved in RNA processing, suggesting that CHD8 occupies a high position in the gene expression control network (Ceballos-Chavez et al., 2015; Subtil-Rodriguez et al., 2014; Sugathan et al., 2014; Wade et al., 2018; Wang et al., 2015; Wang et al., 2017). Furthermore, CHD8 is essential for cell proliferation (Rodriguez-Paredes et al., 2009), since it positively regulates G1/S transition genes during the cell cycle. CHD8 helps to recruit activating E2F proteins to promoters (Subtil-Rodriguez et al., 2014). In addition, CHD8 is necessary for the activation of distal regulatory elements in cis (enhancers) by the hormone progesterone in breast cancer cells (Ceballos-Chavez et al., 2015) and TGFβ-dependent enhancers in neural progenitors (Fueyo et al., 2018).

Development of the central nervous system is very sensitive to a reduction in the level of CHD8. Thus, heterozygous mutations of the *CHD8* gene cause a human syndrome characterized by autism, macrocephaly, distinct facial features and some other clinical characteristics (Barnard et al., 2015; Bernier et al., 2014; Neale et al., 2012; O’Roak et al., 2014; O’Roak et al., 2012a; O’Roak et al., 2012b) (OMIM #610528 and #615032). In fact, CHD8 is one of the strongest autism spectrum disorder (ASD) high-risk associated genes with more than 100 *CHD8* mutations described until now (https://gene.sfari.org/database/human-gene/CHD8). Recently several important works have shown that haploinsufficiency of *Chd8* in mice causes macrocephaly, morphological alterations of cortex and changes in behavior, such as anxiety and alterations in social behaviour (Durak et al., 2016; Gompers et al., 2017; Jung et al., 2018; Katayama et al., 2016), reproducing the symptoms of patients with mutations in *CHD8*. In addition, CHD8 deficiency appears to affect myelination of axons and adipocyte differentiation (Kita et al., 2018; Zhao et al., 2018). All these data demonstrate broad roles of CHD8 in transcription, both in neuronal and non-neural tissues.

How CHD8 is recruited to its target genes is a debated issue and recruitment mediated by RNA polymerase II (RNAPII), other chromatin factors, histone modifications and transcription factors (TFs) have been proposed (Ceballos-Chavez et al., 2015; Dawson et al., 2011; Dou et al., 2005; Kita et al., 2018; Rodriguez-Paredes et al., 2009; Shen et al., 2015; Yates et al., 2010). We have previously described that CHD8 is displaced from its binding sites in several promoters by serum starvation and that serum re-addition triggers the increase of CHD8 occupancy. In this manuscript we demonstrate that the ERK-ELK branch of the mitogen-activated protein kinase (MAPK) pathway is required for serum-dependent recruitment of CHD8 to its target genes. In addition, we show that serum-dependent gene induction, a process mediated by the ERK-ELK pathway, is impaired in CHD8-depleted cells.

## Materials and Methods

### Cell culture and experimental conditions

hTERT-RPE1 (immortalized retina epithelium, RPE1) cell line was maintained in DMEM F12Ham supplemented with 10% fetal bovine serum (FBS), 100 U/ml penicillin and 100 mg/ml streptomycin, in a 37°C incubator, with 5% CO_2_. Cells were grown exponentially (in 10% FBS) and then subjected to serum starvation during 48 h. After that, cells were collected (0%) or medium was replaced with fresh medium supplemented with 20% FBS for the indicated times.

In experiments were the inhibitor of MEK1/2 was used, 50 µM PD98059 (Calbiochem) or vehicle was added to the cells for 1 h, and then medium was replaced by fresh medium supplemented with 20% of FBS for the indicated times.

### RNAi experiments

All siRNAs were transfected using Oligofectamine (Invitrogen) according to the manufacturer’s instructions, with following siRNA sequences: for siCHD8, 5′-GAGCAAGCUCAACACCAUC-3′; siELK1, 5′-GGCAAUGGCCACAUCAUCU-3′; siELK4, 5’-CGACACAGACAUUGAUUCAUU -3’, and siCt, 5′-CGUACGCGGAAUACUUCGA-3′. After transfection, cells were subjected to serum starvation for 48h. After that, medium was replaced by fresh medium with 20% of FBS for the indicated times. The down-regulation of CHD8, ELK1 or ELK4 was confirmed by Western blotting. α-GAPDH antibody (sc-47724) from Santa Cruz Biotechnology was used as a loading control.

### RNA extraction and RT-PCR

Total RNA was prepared by using the RNeasy Kit (Qiagen), as described in the manufacturer’s instructions; note that the step of DNase I digestion was included to avoid potential DNA contamination. cDNA was generated from 1 µg of total RNA using Superscript First Strand Synthesis System (Invitrogen). cDNA (2 μl) was used as a template for qPCR. Gene products were quantified by real-time PCR with the Applied Biosystems 7500 FAST real-time PCR system, using Applied Biosystems Power SYBR green master mix. Sequences of all oligonucleotides are available on Supplementary Table S4. Values were normalized to the expression of the ACTB housekeeping gene. Each experiment was performed at least in duplicate, and qPCR quantifications were performed in triplicate.

### ChIP-qPCR assays

ChIP assays were performed as described (Strutt and Paro, 1999) using anti-CHD8 (A301-224A) from Bethyl Laboratories, anti-ELK1 (ab32106) from Abcam, and anti-ELK4 (H-167) (sc-13030) from Santa Cruz Biotechnology. Chromatin was sonicated to an average fragment size of 400 to 500 bp using the Diagenode Bioruptor. Rabbit IgG (Sigma) was used as a control for non-specific interactions. Input was prepared with 10% of the chromatin material used for immunoprecipation. Input material was diluted 1:10 before PCR amplification. Quantification of immunoprecipitated DNA was performed by real-time PCR (qPCR) with the Applied Biosystems 7500 FAST real-time PCR system, using Applied Biosystems Power SYBR green master mix. Sample quantifications by qPCR were performed in duplicate. Sequences of all oligonucleotides are available on Supplementary Table S4. Data are the average of at least two independent experiments.

### Antibodies and Western blotting

Antibodies used for western blotting were: anti-ELK1 (ab32106) from Abcam, anti-ELK4 (H-167) (sc-13030) and α-GAPDH antibody (sc-47724) from Santa Cruz Biotechnology, anti-CHD8 (A301-224A) from Bethyl, anti-ERK1/2 (ref 4695), P-ERK1/2 (Thr202/-Tyr204, ref 4370), SAPK/JNK (9258), P-SAPK/JNK (Thr202/Tyr204, ref 4668), P38 MAPK (D13E1) (ref. 8690), P-P38 (Thr180/Tyr182, ref 4511) from Cell Signaling Technology and anti-α-tubulin (T9026) from Sigma Aldrich.

### Microarray expression analysis

RPE1 cells were transfected with siControl or siCHD8 and subjected to serum starvation during 48 h. Then, samples for RNA isolation were taken either before or after 1 h of 20% serum stimulation. Total RNA was isolated from three independent experiments by using RNeasy Mini Kit (Qiagen). GeneChip® human Gene 1.0 ST array oligonucleotide microarray (Affymetrix, Santa Clara, CA) were hybridized and scanned as previously described (Rivero et al., 2015). The raw array data were pre-processed and normalized using the Robust Multichip Average (RMA) method (Irizarry et al., 2003). Then differential gene expression analysis was performed using the limma (Ritchie et al., 2015). Genes differentially expressed in CHD8-depleted versus control cells more 1.25 fold (lineal change) and with FDR < 0.05 were selected for further analysis.

### ChIP-seq data analysis

ChIP-seq peaks datasets for T47D (GSE62428), human neural stem cells (GSE57369) or human neuronal progenitor cells (GSE61487) or mouse brain tissues (GSE140117, GSE57369) (Ceballos-Chavez et al., 2015; Cotney et al., 2015; Sugathan et al., 2014; Yildirim et al., 2019) were downloaded from GEO database. Peaks coordinates were centered (peak center +/-500 bp).

Motif binding analyses was performed with CentriMo (v5.0.2) (Bailey and Machanick, 2012) from MEME suite against all 579 binding motifs from JASPAR vertebrates (Mathelier et al., 2015) with default options. CentriMo perform “central motif enrichment analysis” (CMEA), which better identified binding motifs involved in the regulation of ChIP-seq peaks. For each category, CHD8 peaks sequences were shuffled five times for obtaining 5x random sequences with the same %CG content using scrambleFasta.pl from HOMER (v4.10.3) suite, and use that set of sequences as a background. Plots of motif frequency observed versus expected using the background were drown. To estimate significance of the enrichment CentriMo provides the p-value of the one-sided Fisher’s exact test corrected for the number of regions and score thresholds tested (“Multiple Tests”).

ChIP-seq peaks dataset for Elk1 and Chd8 from brain tissue were downloaded from GEO (GSE140117, GSE57369). Overlapping between Chd8 and Elk1 peaks at TSS was determined using bedtools intersect with -u option against all TSS region (TSS +/-1000 bp) for all hg19 UCSC KnownGene transcripts and represented using Venny (https://bioinfogp.cnb.csic.es/tools/venny/). Significance of the overlapping enrichment was determined by using the hypergeometric distribution probability (https://systems.crump.ucla.edu/hypergeometric/index.php).

## Results

### Serum-dependent binding of CHD8 to chromatin is mediated by the MEK1/2-ERK1/2 pathway

We have previously shown that serum-starvation causes the removal of CHD8 from the promoters of CHD8-controled genes, while re-addition of serum promotes the increase of CHD8 occupancy (Subtil-Rodriguez et al., 2014). Then, we decided to investigate which signaling factors are involved in this regulation. The MAPK cascade is one of the main signaling pathways that transmits growth signals to the nucleus. Specially, the MEK1/2-ERK1/2 branch has been extensively involved in the regulation of transcription at several levels, including activation of specific TFs (Gille et al., 1995; Monje et al., 2005), regulation of the general transcriptional machinery (Zhao et al., 2003) and chromatin factors (Courcelles et al., 2013; Goke et al., 2013; Tee et al., 2014). Because of that, we decided to check whether inhibition of the MEK1/2-ERK1/2 pathway affects the recruitment of CHD8 to chromatin after serum addition to serum-starved (quiescent) retinal pigment epithelium, RPE1, cells. RPE1 are non-tumoral human cells with normal karyotype, immortalized by the expression of the telomerase gene (hTERT-RPE1). First, we verified that in our conditions, increasing concentrations of the MEK1/2 inhibitor PD98059 impaired serum-dependent phosphorylation of ERK1/2. A concentration of 50 µM of PD98059 during 1 h prior to serum addition caused about 75% inhibition of ERK1/2 phosphorylation and was used in the following experiments (Fig. 1A). Then, we determined by chromatin immunoprecipitation (ChIP) the kinetics of CHD8 recruitment to the promoters of five CHD8-dependent genes (*CCNA2, CDC6, CCNE2, BRCA2* and *MYC)* (Subtil-Rodriguez et al., 2014) after serum addition to quiescent cells, in the presence or absence of the inhibitor (Fig. 1B). PD98059 treatment significantly impaired CHD8 recruitment to the promoters of the five tested genes (Fig. 1C), suggesting that normal signaling through the MEK1/2-ERK1/2 pathway is required for CHD8 recruitment.

**Fig. 1.**
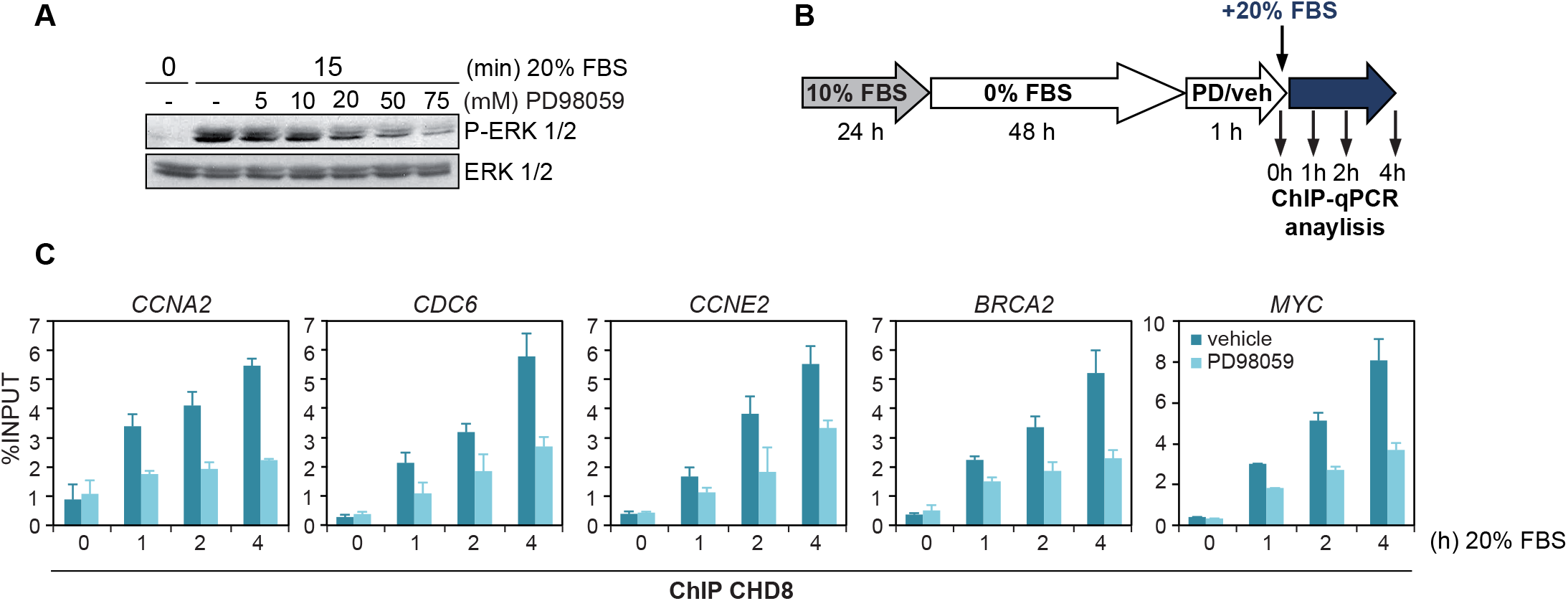
Serum-dependent binding of CHD8 to chromatin is mediated by the MEK1/2-ERK1/2 pathway. A. Western blotting of phospho-ERK1/2 (pERK1/2) 15 minutes after addition of serum (20%) to quiescent RPE1 cells treated with the indicated concentration of PD98059. Level of total ERK1/2 was determined as a control. B. Workflow for the experiment shown in C. PD/veh = PD98059 or vehicle. C. ChIP analysis of CHD8 on selected gene promoters. RPE1 cells were serum-starved for 48 h and then either incubated or not for 1 h with 50 µM of PD98059. Then cells were serum stimulated (FBS 20%) for the indicated times. Data are means ± SD. P < 0.01 for the comparisons of the PD98059 time series versus the vehicle time series, by one-way ANOVA.

### ELK1 and ELK4 binding sites are enriched in CHD8-bound regions

In order to identify TFs that could be responsible for CHD8 recruitment, we performed TF binding motif enrichment analysis from different CHD8 occupancy ChIP-seq data sets, including our previous data and other available data sets (Ceballos-Chavez et al., 2015; Cotney et al., 2015; Sugathan et al., 2014). For that we used CentriMo (Bailey and Machanick, 2012), an algorithm that identifies the region of maximum central enrichment in a set of ChIP-seq peaks and displays the positional distribution of sites. Supplementary Table S1 shows that ETS-like 1 (ELK1) (ACCGGAAGTR) and the related ETS-like 4 (ELK4) (BCRCTTCCGGB) motifs were among the top 10 most enriched motifs in CHD8 peaks in all the studies analyzed, where different antibodies against CHD8 were used in ChIP-seq experiments. ELK1 or ELK4 binding motifs were present in about 50% of the CHD8 peaks found (Supplementary Table S2). Furthermore, CentriMo plots evidenced that ELK1 and ELK4 sites were enriched at the center of CHD8 ChIP-seq peaks (Fig. 2A). Next, we compared the distribution of Chd8 and Elk1 actual occupancy, determined by ChIP-seq at transcription start sites (TSS) from mouse brain tissue. About 75% of the Chd8 ChIP-seq peaks overlap with Elk1 ChIP-seq peaks (Fig. 2B and 2C). Interestingly, about 30% of the Simons Foundation Autism Research Initiative (SFARI) genes (288 out of 954) were bound by both factors in mouse brain which is much more than randomly expected (p = 1.16 × 10^−37^) (Fig. 2D). These data suggest that ELK1 and/or ELK4 factors might cooperate with CHD8. Since ELK1 and ELK4 are well characterized targets of ERK1/2, we hypothesized that these factors may be involved in serum-dependent recruitment of CHD8 to chromatin.

**Fig. 2.**
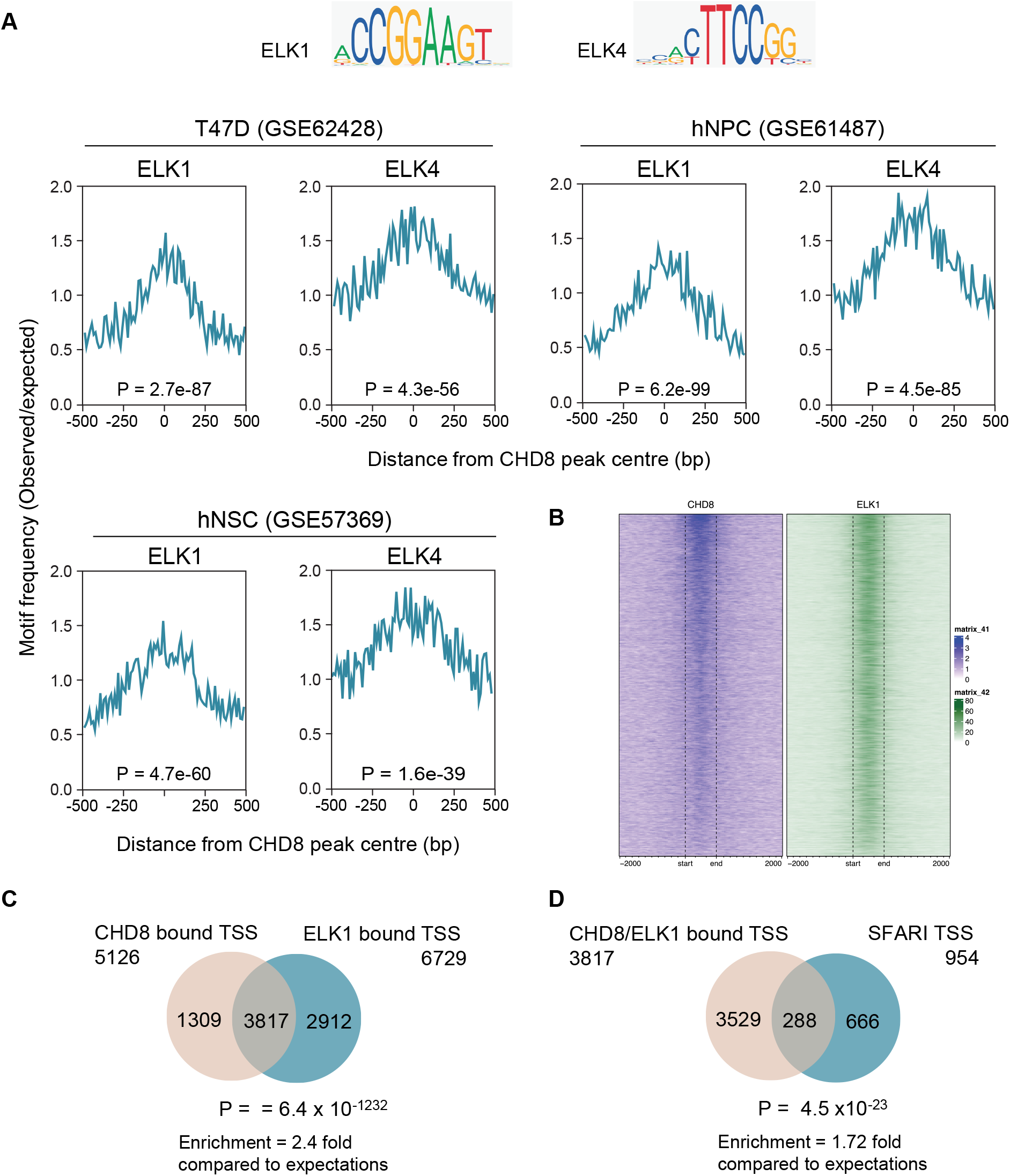
ELK1 and ELK4 DNA binding sites are enriched in CHD8-bound regions. A. Frequency of ELK1 and ELK4 motifs in CHD8 peaks from the indicated datasets. Observed versus expected frequencies are represented. Expected frequencies were calculated from random background constructed as described in Materials and Methods. P-value of the enrichment using Fisher exact test is provided. B. Heat map showing occupancy of Chd8 and Elk1 in mouse brain (ChIP-seq signal from GSE140117 and GSE57369). C. Venn diagram showing overlapping of TSS that present CHD8 and ELK1 determined by ChIP-seq. D. Venn diagram showing overlapping of CHD8 and ELK1 containing TSS with ASD-associated genes as reported by SFARI (https://gene.sfari.org). C, D. Significance of the overlapping enrichment using hypergeometric distribution probability is provided.

### ELK1 and ELK4 contributes to CHD8 recruitment to chromatin

To verify whether ELK1 mediates recruitment of CHD8, we analyzed serum-dependent binding of ELK1 and CHD8 to several target promoters by ChIP-qPCR, in cells depleted of ELK1 or CHD8 by RNA interference. For that, cells were transfected with siRNAs, then subjected to serum starvation for 48 h and then serum stimulated. Samples for ChIP-qPCR analysis were taken under quiescent conditions or 1 and 4 h after adding serum to quiescent cells (Fig. 3A). Depletion of ELK1 and CHD8 proteins was verified by western blotting (Fig. 3B). First we verified that serum stimulation of quiescent cells increased occupancy of ELK1 (Fig. 3C) at the *CCNA2, CDC6, CCNE2, BRCA2* and *MYC* promoters. ELK1 depletion strongly decreased the ChIP signals, validating the specificity of the antibody. Interestingly, depletion of CHD8 did not significantly affect ELK1 occupancy after serum re-addition (Fig. 3C), indicating that ELK1 recruitment is independent of CHD8. However, depletion of ELK1 significantly decreased occupancy of CHD8 (Fig. 3D). As a control, we verified that silencing CHD8 strongly decreased CHD8 ChIP signals validating the specificity of the antibody (Fig. 3D). Next, we examined whether ELK4 is also involved in serum-dependent CHD8 recruitment to chromatin. First, we demonstrated by ChIP-qPCR that ELK4 protein binds the promoters of the selected genes and that, as shown for ELK1, binding is independent of CHD8 (Fig. 3E). Then, we showed that depletion of ELK4 significantly impairs CHD8 recruiting as well (Fig. 3E). Efficiency of ELK4 depletion by the corresponding siRNAs was verified by western blot (Fig. 3B).

**Fig. 3.**
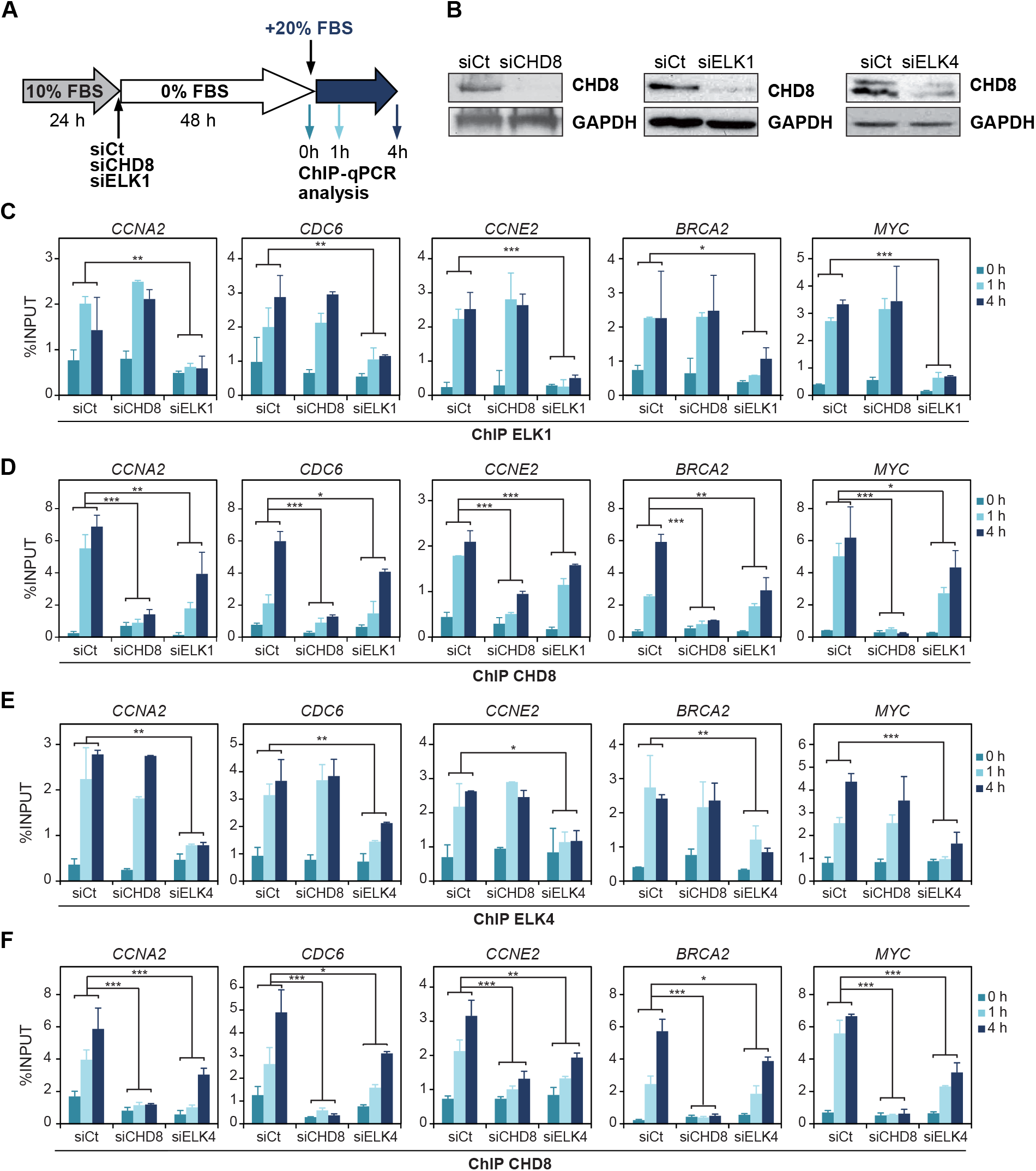
ELK1 and ELK4 contribute to serum-dependent recruitment of CHD8 to chromatin. A. Workflow for the experiment shown in C and F. B. Efficiency of CHD8, ELK1 and ELK4 knockdown was determined by western blotting. C-F. ChIP analysis of ELK1 (C), CHD8 (D, F) and ELK4 (E) at the indicated promoters in RPE1 cells serum starved and then stimulated with 20% FBS. Cells were transfected with the indicated siRNA. Data (% input) are the mean of at least two experiments, where three independent time points (0h, 1h and 4h) were taken per each experiment. Error bars represent ± SD values. A two-ways ANOVA was used for statistical analysis. *p < 0.05, **p < 0.01 versus control.

Functional redundancy between ELK1 and ELK4 has been demonstrated using mouse models. Thus, Elk4(-/-) and Elk1(-/-) mouse single mutants show minimal phenotypic changes, however, double Elk4(-/-) Elk1(-/-) animals present several phenotypes including defects in T-cell differentiation (Costello et al., 2010; Maurice et al., 2018). Our data indicate that ELK1 and ELK4 are not fully redundant for CHD8 recruitment, since silencing of each one of the factors has phenotypic consequences in CHD8 occupancy. Interestingly, depletion of both factors promoted similar impairment of CHD8 occupancy than depletion of each factor separately (Fig. 4). In addition, these data demonstrated that a residual binding of CHD8 to the chromatin occurs even upon depletion of both factors. Although, this can be a consequence of partial silencing of ELK factors, we cannot rule out that other additional factors are involved in CHD8 recruitment.

**Fig. 4.**
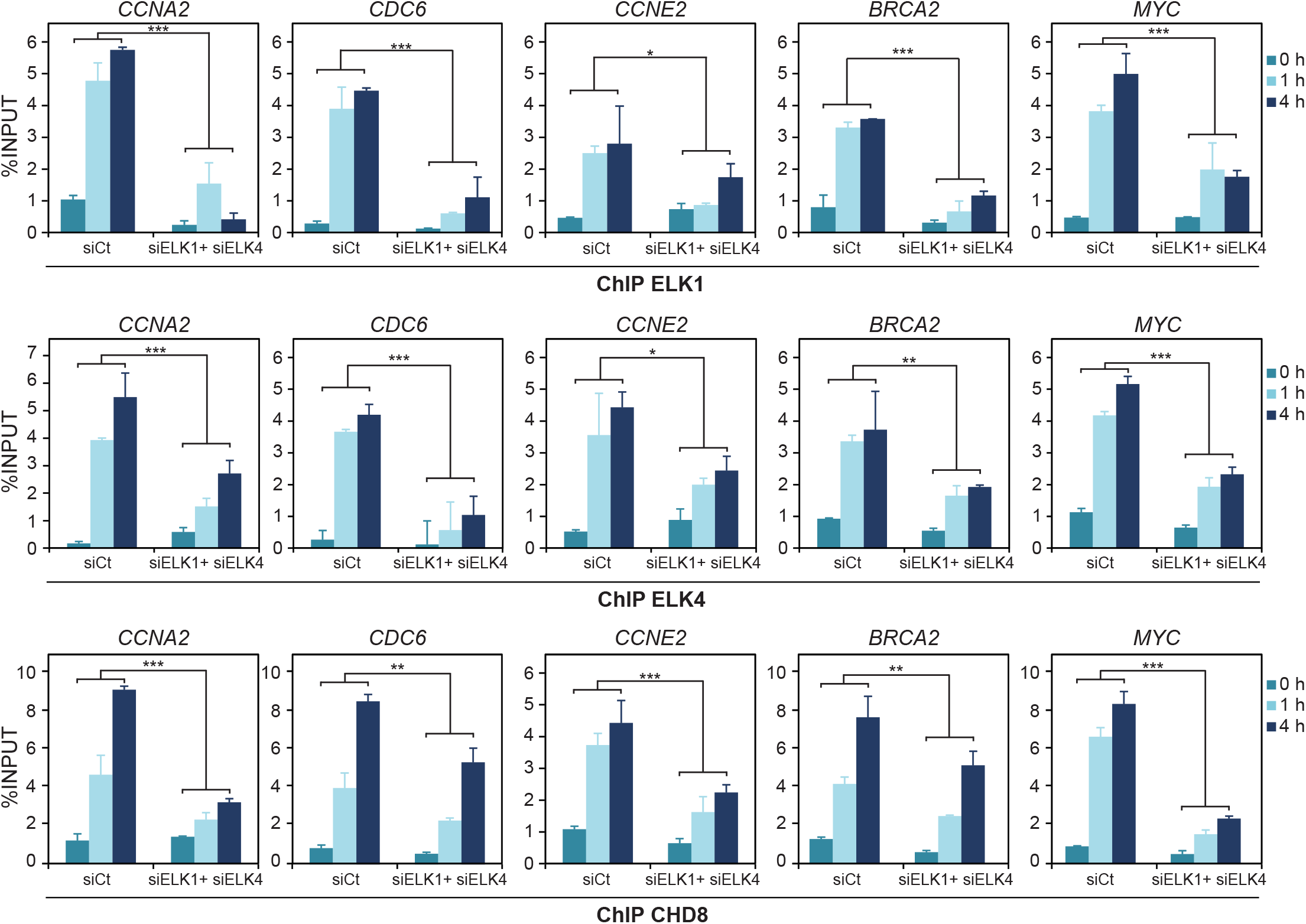
A fraction of chromatin-associated CHD8 is independent of ELK1 and ELK4. ChIP analysis of ELK1, ELK4 and CHD8 at the indicated promoters, in RPE1 cells serum starved and then stimulated with 20% FBS. Cells were transfected with control siRNA (siCt) or siRNA against ELK1 and ELK4 at the same time (siELK1siELK4). Data (% input) are the mean of at least two experiments were three independent time points (0h, 1h and 4h) were taken per each experiment. Error bars represent ± SD values. A two-ways ANOVA was used for statistical analysis. *p < 0.05, **p < 0.01, ***p < 0.001 versus control.

### CHD8 controls serum-dependent activation of gene expression

It is well known that the ERK-ELK pathway controls serum-dependent activation of gene expression together with the SRF transcription factor (Esnault et al., 2017; Hashimoto et al., 2011; Hipskind et al., 1994; Latinkic et al., 1996; Wozniak et al., 2012). Since we have shown that CHD8 is recruited to specific promoters by ELK1 and ELK4, we wondered whether gene induction by serum, at a whole-genome level, was dependent on CHD8. To test this hypothesis, we analyzed the changes in the transcriptome one hour upon serum stimulation of quiescent RPE1 cells transfected with a control siRNA (siCt) or a siRNA targeting CHD8 (siCHD8) (Fig. 5A). 583 genes were upregulated by serum (Fold change >1.25 and FDR<0.05) in control cells, including typical immediate-early genes such as *FOS, FOSB, JUN, EGR2* and *MYC* (Supplementary Table S3). Importantly, induction of expression of a subset of genes was impaired in CHD8-depleted cells (Fig. 5B and Supplementary Table S3). Thus, 46.0% of the serum-induced genes were less activated in CHD8-depleted cells than in control cells (less than 80% activation with respect to siCt) (Fig. 5C and Supplementary Table S3). 46.5% and 7.5% genes were not affected and more activated (more than 125% activation), respectively, in CHD8-depleted cells compared with control cells. CHD8 dependence was confirmed by RT-qPCR in three well-known serum-induced genes *JUN, FOS* and *MYC* (Fig. 5D). Interestingly, *FOS* and *MYC* are known targets of ELK1 (Hashimoto et al., 2011; Koenig et al., 2010; Price et al., 1995) (and Fig. 3). As a control, we verified that depletion of CHD8 does not significantly impairs activation of the more relevant MAPK pathways (ERK1/2, JNK and p38) (Supplementary Figure S1), suggesting that impaired activation of serum responsive genes in CHD8-depleted cells is not a consequence of a deficient signal transduction. Therefore, our data show that CHD8 is required for normal induction of a subset of serum-induced genes.

**Fig. 5.**
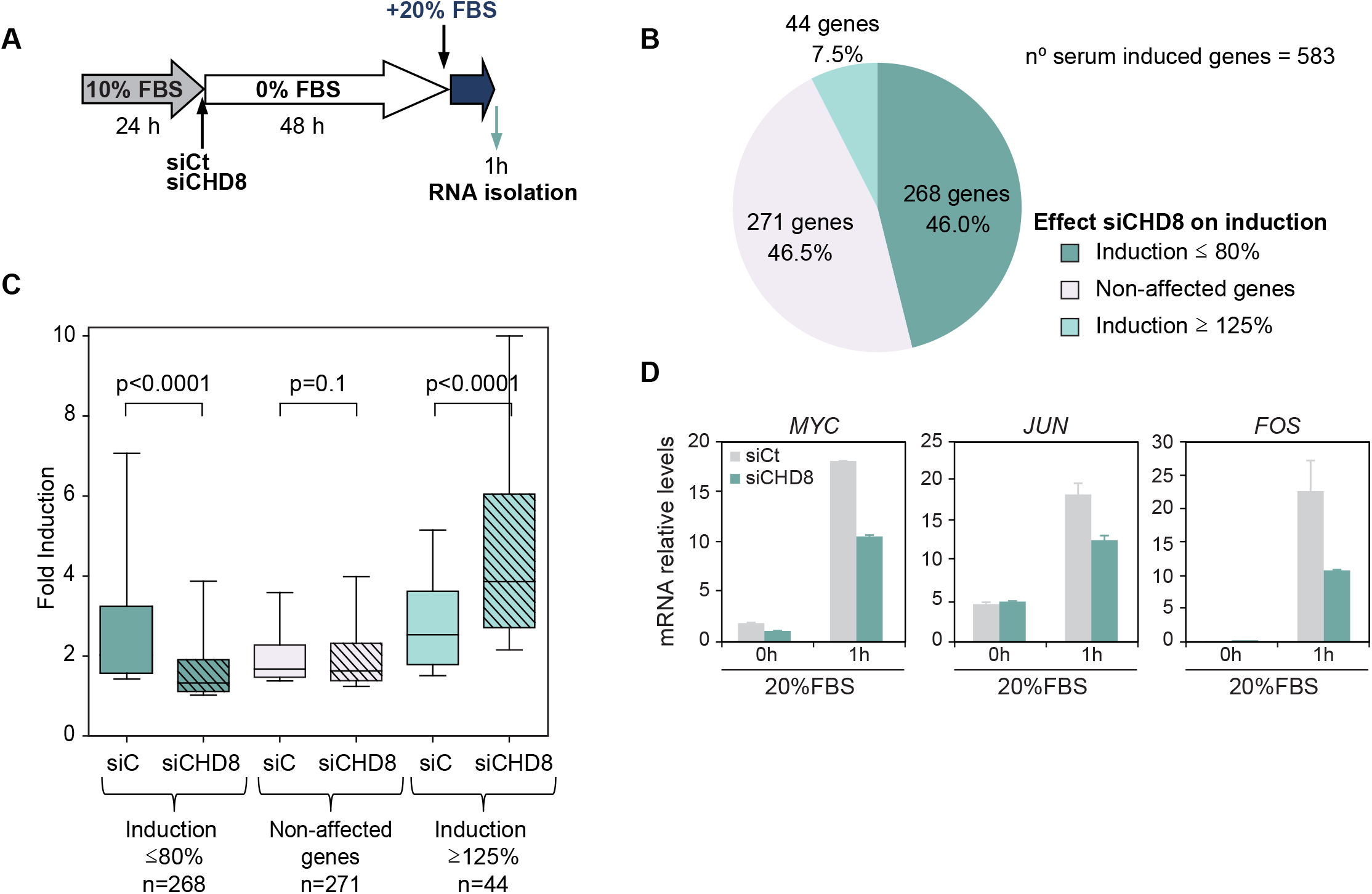
Transcriptomic analysis of serum-induced genes under CHD8 depletion. A. Workflow for the transcriptomic experiment. B. Serum-induced genes were divided into three categories depending on the effect of CHD8 depletion on serum-dependent induction levels: genes induced less than 80%, genes induced more than 125% and non-affected genes. C. Box plot showing fold induction of the different categories of genes classified with respect to its CHD8-dependence. D. RT-qPCR analysis of expression of selected CHD8-dependent genes in RPE1 cells transfected with control siRNA (siCt) or siRNA against CHD8 (siCHD8). Cells were serum starved and then serum stimulated as in (A). mRNA (relative levels) are the mean of three independent experiments. Error bars represent ± SD values.

## Discussion

In the present manuscript we show that the ERK-ELK pathway is essential for correct serum-dependent recruitment to chromatin of CHD8 at five different promoters in non-neuronal cells. We also show that CHD8 is required for the correct induction of a subset of serum-induced genes, including typical ERK-ELK targets. Interestingly, a recent preprint also demonstrates a role of ELK1 in CHD8 recruiting in mouse neurons, although effect of serum was not evaluated in their experiments.

We show that ELK1 and ELK4 are required for CHD8 recruitment to promoters, but CHD8 is not necessary for ELK1 or ELK4 binding. Consistent with our data, recent results demonstrate that ELK proteins are required for many histone modifications at serum-dependent promoters, including H3K9acS10ph, H4K16ac, H3K27ac, H3K9acK14ac, and H3K4me3, suggesting that ELK factors binding to the promoters is an early event during the process of promoter activation (Esnault et al., 2017). How are ELK factors involved in CHD8 recruitment? We have tried to demonstrate a direct physical interaction between CHD8 and ELK1 or ELK4 by immunoprecipitation of endogenous proteins but we have not been successful. Ternary complex factors (TCF) (ELK1, ELK4, and ELK3) stabilizes Serum Response Factor (SRF) binding to the DNA and in fact, most SRF genomic binding sites are co-occupied by TCF factors (Gualdrini et al., 2016). A direct interaction between CHD8 and SRF was reported more than 10 years ago (Rodenberg et al., 2010). Therefore, CHD8 recruitment to chromatin might require the cooperation of ELK1 and/or ELK4 with SRF.

CHD8 is one of the top risk genetic factors that causes ASD (Barnard et al., 2015; Bernier et al., 2014; Neale et al., 2012; O’Roak et al., 2014; O’Roak et al., 2012a; O’Roak et al., 2012b). To our knowledge, ELK1 and ELK4 have not yet been associated with ASD. However, it is well known that the ERK signaling pathway is critical for brain development (Samuels et al., 2009). In the developing cortex, the ERK pathway is essential for neuronal responses to neurotransmitters and receptor tyrosine kinase ligands. In fact, deficiencies in ERK signaling during development have been associated with ASD (Levitt and Campbell, 2009; Xing et al., 2016; Yufune et al., 2015). Furthermore, the ERK-ELK pathway plays an essential role in neuronal activity-dependent immediate-early genes expression (Shilyansky et al., 2010; Thomas and Huganir, 2004; Tyssowski et al., 2018). Interestingly, serum-dependent induction of eight genes that have been shown to be induced by neuronal activity (Ataman et al., 2016; Madabhushi et al., 2015) was impaired upon knockdown of CHD8 in our system (*FOS, FOSB, PTGS2, NR4A1, EGR2, NR4A2, NR4A3 and NPAS4*). All these data point to a role of the ERK-ELK-CHD8 axis in neuronal function and eventually in ASD etiology.

## Availability of data and materials

Chip-seq data used in this manuscript can be accessed at the Gene Expression Omnibus (GEO), (https://www.ncbi.nlm.nih.gov/geo/), accession numbers. GSE62428, GSE57369, GSE61487, GSE91713, GSE140117. All data generated are included in this published article and additional files.

## Funding

This work was funded by the Spanish Ministry of Economy and Competitiveness MCIN/AEI/10.13039/501100011033/ (BFU2017-85420-R and PID2020-118516GB-I00), the Junta de Andalucía (BIO-321 and Proyecto PAIDI2020 PY18-1962), and the European Union (FEDER). CABIMER is a Center partially funded by the Junta de Andalucía.

## Authors’ contributions

AS-R and JCR, conceived the project. AS-R and EV-C, did major experiments. analyzed the data. JAG-M and MC-C, analyzed data. JCR and MC-C, wrote the manuscript.

## Acknowledgements

We are grateful to M. García-Domínguez for critical reading of the manuscript and discussion. We thank E. Andújar and M. Pérez from the CABIMER Genomic Unit for microarray expression analysis.

## Competing interests

The authors declare that they have no competing interests.

## Supplementary data legends

**Supplementary Figure S1. Effect of CHD8 depletion on MAPK signaling**. RPE1 cells were transfected with control siRNA (siCt) or siRNA against CHD8 (siCHD8), subjected to 48 h of serum starvation and then, serum stimulated (20% FBS). Samples for western blotting were taken at the indicated times.

**Supplementary Table S1**. Analysis of enriched DNA binding motifs at CHD8 ChIP-seq peaks from different datasets using Centrimo. ChiP-seq data from neuronal stem cells using antibody ab114126 from Abcam (GSE57369); neuronal progenitors using antibodies A301-224A (Bethyl), NB100-60417 (Novus), and NB100-60418 (Novus) (GSE61487) and T47D breast cancer cells using antibody A301-224A (Bethyl) (GSE62428).

**Supplementary Table S2**. Percentage of CHD8 peaks from different ChIP-seq experiments, containing the indicated binding motifs. Data obtained using CentriMo analysis.

**Supplementary Table S3**. Genes upregulated 1 h after serum addition to serum-starved RPE1 cells transfected either with siControl (siCt) or siCHD8.

**Supplementary Table S4**. Oligonucleotides used in this study.

## Notes

### Competing Interest Statement

The authors have declared no competing interest.

