## Supplementary figures and images for "Serum-dependent recruitment of the chromatin remodeler CHD8 to promoters is mediated by the ERK-ELK pathway"

### Supplementary Figure S1

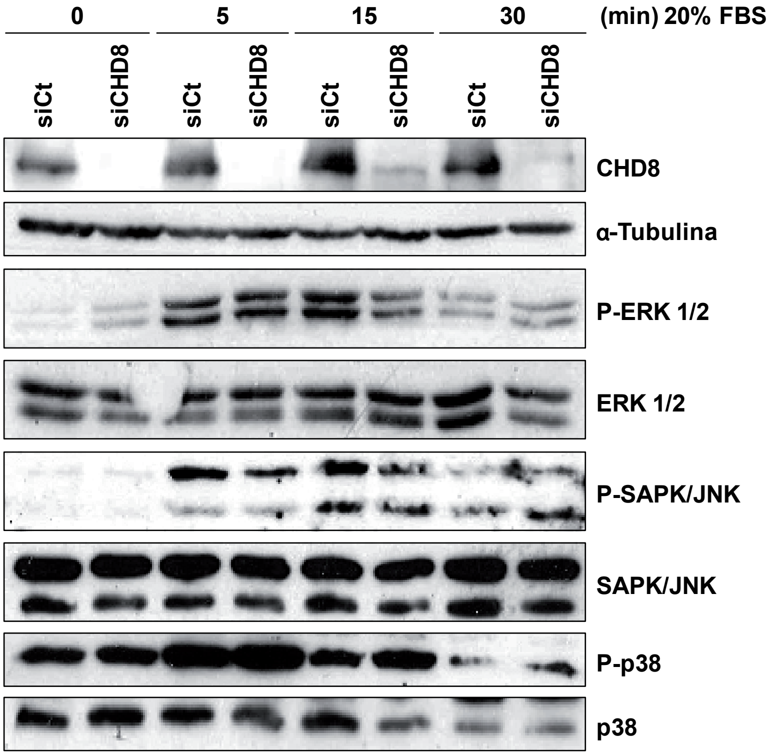


Supplementary figure 1
