## Supplementary Table S2 for "Serum-dependent recruitment of the chromatin remodeler CHD8 to promoters is mediated by the ERK-ELK pathway"

Supplementary Table S2. Percentage of CHD8 peaks from different ChIP-seq experiments, containing the indicated binding motifs. Data obtained using CentriMo analysis.

| | Cotney et al., 2015 (GSE57369) | Sugathan et al., 2014 (GSE61487) | | | Ceballos-Chávez et al 2014. (GSE62428) |
|---|---|---|---|---|---|
| Binding motif | Abcam ab114126[a] | Bethyl A301-224A [a] | Novus NB100-60417 [a] | Novus NB100-60418 [a] | Bethyl A301-224A [a] |
| ELK1 | 31% | 32% | 30% | 37% | 26% |
| ELK4 | 51% | 44% | 45% | 42% | 37% |
| ELK1 or ELK4 | 56% | 52% | 51% | 55% | 42% |
| ELK1 and ELK4 | 26% | 25% | 24% | 24% | 21% |

[a]Antibody used in the ChIP-seq experiment.
