## Supplementary Table S4 for "Serum-dependent recruitment of the chromatin remodeler CHD8 to promoters is mediated by the ERK-ELK pathway"

Supplementary Table S4. Oligonucleotides used in this study.

| Loci | Forward 5’-3’ | Reverse 5’-3’ | Used for |
|---|---|---|---|
| *CCNA2* | CAGTGCCTGGGGTTTAAAAG | GGGGTCAAACCAAGCTCTAA | ChIP |
| *CDC6* | GTAAAAGCCCTGCCTCTCAG | ATGGATGGTTTACCCAGACG | ChIP |
| *CCNE2* | TGACACCCCCGAAATCCA | CCTGGCTCGCGCATCT | ChIP |
| *BRCA2* | AAGCGTGAGGGGACAGATT | GCCGGAGTAAGCTGACAAA | ChIP |
| *MYC* | GCCTGGAGGCAGGAGTAAT | GTAGCTTCCAAATCCGATGC | ChIP |
| *MYC* | GCCTGGAGGCAGGAGTAAT | GTAGCTTCCAAATCCGATGC | RT-qPCR |
| *JUN* | TGACTGCAAAGATGGAAACG | CAGGGTCATGCTCTGTTTCA | RT-qPCR |
| *FOS* | CGAGCGCAGAGCATTGG | CCTTCGGATTCTCCTTTTCTCTT | RT-qPCR |
| *ACTB* | ACGAGGCCCAGAGCAAGA | GACGATGCCGTGCTCGAT | RT-qPCR |
